# LIRcentral: a manually curated online database of experimentally validated functional LIR-motifs

**DOI:** 10.1101/2022.06.21.496832

**Authors:** Agathangelos Chatzichristofi, Vasileios Sagris, Aristos Pallaris, Marios Eftychiou, Ioanna Kalvari, Nicholas Price, Theodosios Theodosiou, Ioannis Iliopoulos, Ioannis P. Nezis, Vasilis J Promponas

## Abstract

Several selective macroautophagy receptor and adaptor proteins bind members of the Autophagy-related protein 8 (Atg8) family using short linear motifs (SLiMs), most often referred to as Atg8-interacting motifs (AIM) or LC3-interacting motifs (LIR). AIM/LIR-motifs have been extensively studied during the last fifteen years, since they can uncover the underlying biological mechanisms and possible substrates for this key catabolic process of eukaryotic cells. Prompted by the fact that experimental information regarding LIR-motifs can be found scattered across heterogeneous literature resources, we have developed LIRcentral (https://lircentral.eu), a freely available online repository for user-friendly access to comprehensive, high-quality information regarding LIR-motifs from manually curated publications. Herein, we describe the development of LIRcentral and showcase currently available data and features, along with our plans for the expansion of this resource. Information incorporated in LIRcentral is useful for accomplishing a variety of research tasks, including: (i) guiding wet biology researchers for the characterization of novel instances of LIR-motifs, (ii) giving bioinformaticians/computational biologists access to high-quality LIR-motifs for building novel prediction methods for LIR-motifs and LIR containing proteins (LIRCPs) and (iii) performing analyses to better understand the biological importance/features of functional LIR-motifs. We welcome feedback on the LIRcentral content and functionality by all interested researchers and anticipate this work to spearhead a community effort for sustaining this resource which will further promote progress in studying LIR-motifs/LIRCPs.

## Introduction

Autophagy comprises lysosome-mediated degradation processes for recycling cytoplasmic material and plays a central role in eukaryotic cell homeostasis and stress response [1]. Macroautophagy is the best studied form of autophagy; it relies on the nucleation, elongation and maturation of autophagosomes –double-membraned structures engulfing cytoplasmic material to be degraded– which eventually fuse with lysosomes for recycling to occur [2]. Key components of the macroautophagy machinery are members of the Atg8 protein family (MAP1LC3/LC3 and GABARAP in human), small proteins with a ubiquitin-like fold: they participate essentially in all steps of macroautophagy from phagophore nucleation, to autolysosome formation [3]. They are involved in an increasing number of biological processes, including LC3-associated phagocytosis [4, 5], endosomal microautophagy [5, 6], and unconventional secretion [7].

In selective macroautophagy, selectivity is mainly mediated through receptor proteins that bind to the surface of Atg8s by anchoring a small linear peptide (namely the Atg8 interacting motif (AIM) in plants and fungi and the LC3 interacting region motif (LIR-motif) in mammals [8]). Following the discovery of the first few LIR-containing proteins (LIRCPs)[9–11], several definitions of the LIR-motif have appeared in the literature [9,12–15], highlighting a short core consensus sequence described by the regular expression pattern [WFY]xx[VLI].

The collection of a few dozen experimentally characterized LIR-motifs made it possible to develop computational methods for their prediction [16, 17]. Such prediction tools have inevitably facilitated the advancement of research towards identifying the molecular mechanisms underlying several subtypes of selective autophagy and related biological processes [18], and the number of scientific publications reporting the experimental validation of functional LIR-motifs continues to accumulate. Despite the importance of experimentally verified LIR-motifs, for wet lab biologists and computational biologists alike, there is a lack of a central repository of LIR-motif instances curated from the literature describing such sequences across species.

Currently, researchers wishing to obtain information about LIR-motifs have access to online prediction tools (i.e. iLIR [16] and hfAIM [17]) as well as the recently described pLIRm software [19], a new computational tool for discriminating functional LIR-motifs, which is provided as open source Python code. In addition, there exist dedicated databases of predicted LIR-motifs for particular species (i.e. iLIR database [20] and iLIR@viral [21]) which are easy for users to navigate and retrieve relevant predictions.

In the above mentioned databases, the available LIR-motifs are taxonomically restricted (e.g. the iLIR database provides predictions for 8 model species only), thus for species not covered by these resources, interested users need to directly use the available prediction tools. An additional source of data relevant to LIR-motifs has become available with the most recent version of the ELM database of Eukaryotic Linear Motifs [22] and its associated resource articles.ELM [23].

Summarizing, despite the increasing availability of in silico resources for LIR-motifs, a number of inherent limitations exist. With respect to prediction tools, all currently available methods are unable to detect atypical cases of LIR-motifs (e.g. non-canonical LIR-motifs), while pLIRm is only available through the command line, thus preventing easy use by wet biologists. Moreover, the small size of available datasets has hindered the extensive validation of these prediction methods: in lieu of extended datasets of experimentally characterized LIR-motifs, all prediction methods are not extensively/independently validated and it is possible that they suffer from overfitting [19]. On the other hand, relevant databases come with their drawbacks too: iLIR database/iLIR@viral provide predicted LIR-motifs only and in a taxonomically restricted manner, while ELM provides experimentally verified instances of LIR-motifs but with small coverage only (67 LIR-motif instances under four different ELM classes: ELME000368-ELME000371; last accessed May 15, 2022).

An alternative –yet more cumbersome– route to obtain LIR-motif related information supported by experimental evidence is to directly search within the PubMed database [24] or other literature databases, e.g. Europe PMC [25], PubMedCentral [26]. This approach suffers from the fact that relevant information is scattered in a non-standardised manner within hundreds of publications, thus making it difficult to extract relevant data. For example, a PubMed search with the keyword-based query

`‘autophagy AND (“lir motif” OR “aim motif” OR “lir-motif” OR “aim-motif”)’`

(hereinafter ‘LIRquery’) identifies 99 entries (query last performed on May 15, 2022) (**Table S1**). Of these, only 60 (60.7%) present primary characterization data of at least one LIR-motif while the remaining 39 (39.3%) are review papers or editorials on relevant topics. While reviews are unquestionably important in summarizing the literature in any particular topic, they more often contain high-level information and readers wanting to obtain detailed information on particular LIR-motifs (e.g. their originating sequences or their exact positions therein) need to eventually consult the primary research paper that describes the characterization of the motif.

According to our experience, several papers reporting the experimental validation of LIR-motifs often deal with broader topics, thus motif characterization and relevant information may not be directly reflected in the paper title or abstract, which is where the PubMed search engine is searching for the query keywords. When performing the ‘LIRquery’ against the full-text literature databases EuropePMC and PubMedCentral we retrieved 899 and 1150 entries respectively (queries performed on May 24, 2022). In such long lists, it is expected that several papers will only be marginally relevant to the topic of interest (e.g. reviews); many more are completely irrelevant for finding information confirming the functionality of particular LIR-motifs, with a range of cases where LIR-motis are thoroughly reviewed or simply mentioned. For instance, when examining the 20 most recent entries retrieved from EuropePMC with the ‘LIRquery’ only 3 entries (15%) correspond to papers describing the characterization of LIR-motifs (**Table S2**). While this dataset can by no means be considered representative of the underlying body of literature, it gives a first glimpse of the number of false positives returned by similar full-text queries. Carefully studying the text of these literature entries (and often extensive supplementary files) to extract information about LIR-motifs requires tremendous effort, even from experienced researchers. A collection of literature-supported evidence for functional LIR-motifs should provide interested researchers with easy and quick access to data, available in a standardized form, thus facilitating additional research efforts for deciphering the mechanisms of selective macroautophagy and related processes.

Herein, we describe our efforts on the creation of a comprehensive collection of experimentally characterized LIR-motifs based on extensive biomedical literature search, aided by simple text analysis techniques, and followed by meticulous manual curation of full text papers. Furthermore, we present the development of a web accessible database (LIRcentral; https://lircentral.eu) in which interested researchers are able to retrieve information about literature-validated LIR-motifs and display them along with features annotated in UniProt [27]. By cross-referencing protein entities to UniProt entries, LIRcentral enables seamless data integration with other resources. We also provide example cases where using LIRcentral can enable posing and answering interesting questions related to LIR-motifs and LIRCPs in reasonable time, even for users with limited computational skills/background.

## Results and Discussion

### Content of the LIRcentral Database

#### Curated publications in LIRcentral

As of writing this text (June 12, 2022) a total of 159 peer-reviewed manuscripts from 56 different journals (**Table 1**) have served for the curation of LIRcentral entries (See Materials and Methods). Even though information regarding LIR-motifs increasingly often appears in pre-prints or other unreviewed sources of information, we deliberately do not include these motifs in LIRcentral. Instead, we mark LIR-motifs reported in such sources as candidate entries and wait until the work appears in a journal after peer-review for inclusion in LIRcentral. Thus, entries currently in LIRcentral are supported by at least one primary research paper which is manually curated for each LIR-motif. Review papers serve both as an additional means to track the primary literature as well as to resolve any ambiguities therein.

**Table 1.** Sources of journal papers (n=159) reporting the experimental validation of LIR-motifs currently curated for LIRcentral entries.

| **Journal title** | **# papers** | **Journal title** | **# papers** |
| --- | --- | --- | --- |
| Autophagy | 24 | Cell | 2 |
| The Journal of Biological Chemistry | 14 | Cell Death & Disease | 2 |
| Nature Communications | 12 | Cell Metabolism | 2 |
| eLife | 9 | Cell Reports | 2 |
| The Journal of Cell Biology | 9 | Communications Biology | 2 |
| Molecular Cell | 8 | Current Biology | 2 |
| Developmental Cell | 6 | FASEB Journal | 2 |
| EMBO Reports | 5 | PloS One | 2 |
| Nature Cell Biology | 5 | PNAS | 2 |
| Nature | 4 | Science Signaling | 2 |
| Cell Death and Differentiation | 3 | The Plant Cell | 2 |
| Oncotarget | 3 | Other (Journals with 1 paper each) | 32 |
| The EMBO Journal | 3 |  |  |

Papers describing LIR-motifs get published in both niche and more generic journals. More than half of the total number of papers is published in the top-7 journals (in terms of relevant papers published) in this list.

#### Properties of the LIRcentral literature corpus

We attempt to qualitatively compare the content of the abstracts of these 159 documents in contrast to a background composed from the abstracts of 1,000 manuscripts retrieved from PubMed using the keyword ‘autophagy’. By constructing word-clouds, some differences in the prevailing terms in these two (not necessarily disjoint sets of texts) are evident: the manuscripts used to annotate LIR-motif entries are enriched in terms such as ‘binding’, ‘protein’, ‘LC3’, ‘Atg8’, ‘interaction’, ‘selective’, which reflect the role of the LC3::LIR-motif interaction in selective autophagy, while the background set is enriched in more generic terms, e.g. ‘induced’, ‘expression’, ‘treatment’, ‘apoptosis’ (**Figure 1**).

**Figure 1.**
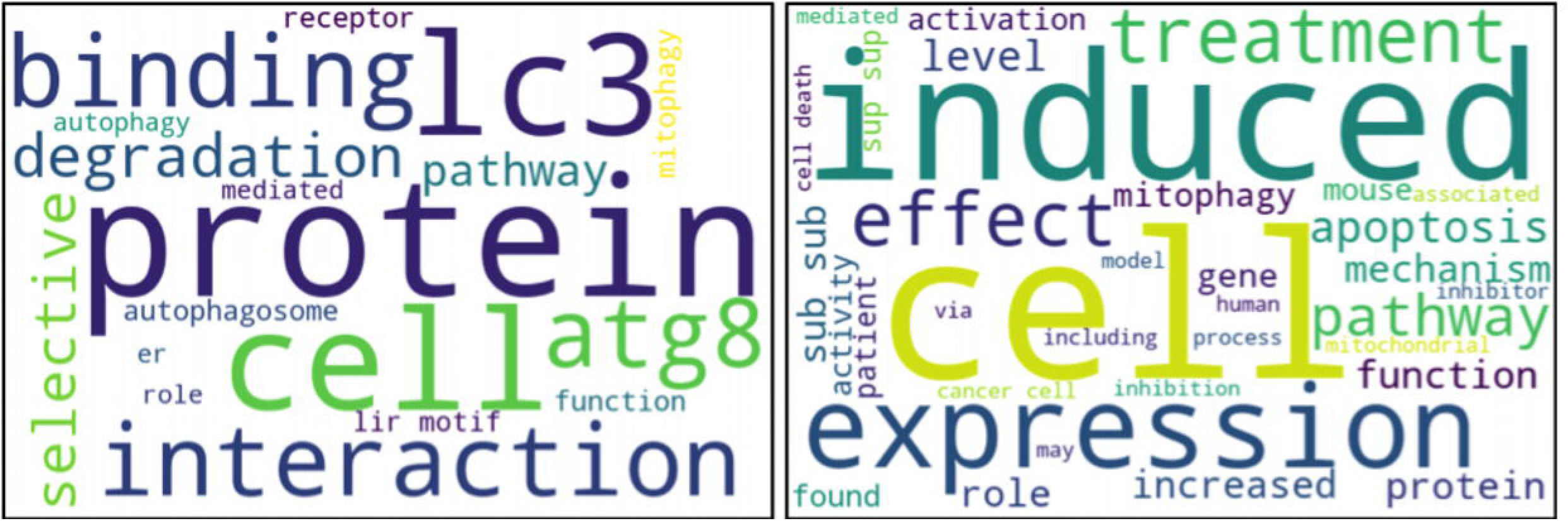
Word cloud representation of the textual content of all abstracts from the papers curated in the current LIRcentral version (left) versus 1000 articles retrieved by the keyword “autophagy” using the Bio:Entrez module of Biopython (right).

#### Comparison of the LIRcentral literature set to other (curated) collections

We compare the sets of curated papers used to populate LIRcentral to other relevant sets of publications: (i) the dataset of publications associated with the LIR-motif instances in the ELM database (n=57; last accessed May 09, 2022)[22], (ii) the list of papers listed to be relevant to LIR-motifs within the articles.ELM resource (n=103; last accessed May 09, 2022)[23], and (iii) the manuscripts linked to instances of LIR-motifs used to train/validate pLIRm (n=46; taken from Supplementary Data 1 in [19]). We also used the results of the LIRquery against PubMed (accessed May 09, 2022) as a control.

From the Venn diagram representation (**Figure 2**) it is obvious that these collections are largely heterogeneous. Only 1 paper is common across all 5 collections: this is the seminal paper on the topic by Alemu and colleagues [15].

**Figure 2.**
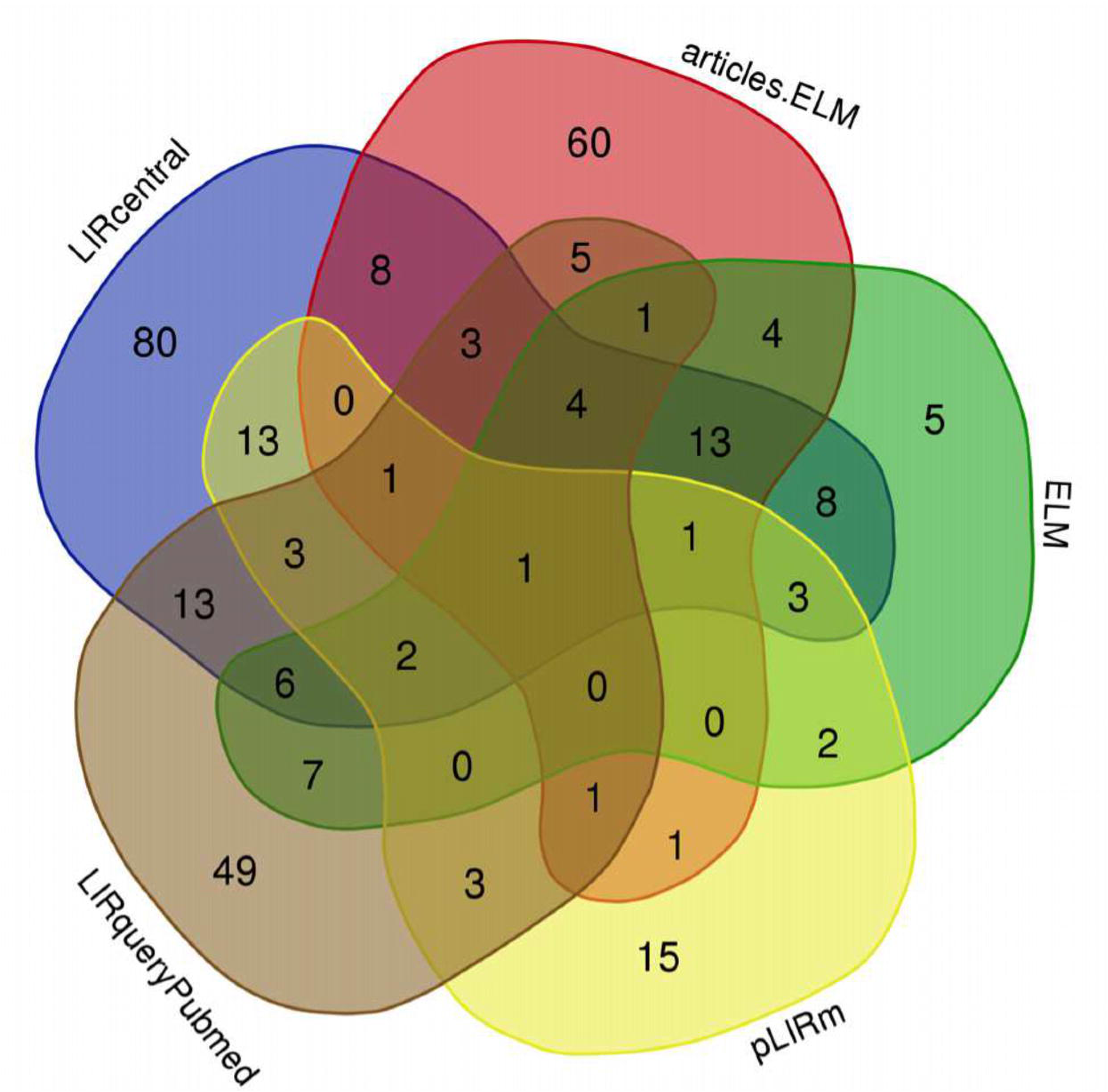
A Venn diagram representation of the overlaps between different literature collections related to LIR-motifs.Drawn using https://bioinformatics.psb.ugent.be/webtools/Venn/.

The LIRcentral literature collection is the most comprehensive and contains the most unique elements (n=80) not available in any other literature dataset. These 80 papers, uniquely available through LIRcentral, span from 2009-2020 and mainly report primary results (only 6 marked as ‘Review’ in PubMed). On the other hand, a total of 153 papers in this collective literature corpus are not curated in the current LIRcentral version of which (i) a large fraction (58, ∼38%) are published during the last 3 years, and (ii) 21 (∼14%) are marked in PubMed as review papers.

The articles.ELM resource contains a large number of unique elements (n=60) several of which are either false positive predictions – i.e. they do not describe experimental validation of LIR-motifs– (e.g. papers with PubMed identifiers (PMIDs): 30078701, 28388439, 27422009, 27594685) or deal with other important functional aspects of LIR-motifs, including post-translational modifications (e.g. PMID: 27757847) or the higher order structural organization and interactions of LIRCPs (e.g., PMID: 25921531, 26506893); however, it is worth noting that this set has been assessed without taking into account the confidence score accompanying each publication.

Even though we have not manually checked all papers in detail, from the 5 unique ELM literature citations two were apparently unrelated to LIR-motifs (PMID: 24668265, 24668266). These were used for annotating motif instance ELMI004471 in human WD repeat and FYVE domain-containing protein 3 (Alfy; UniProt: Q8IZQ1) and were communicated to the ELM curators for correction (June 11, 2022).

There were also 15 papers appearing in the pLIRm collection only. These were published between 2014 and 2020, with 3 entries marked by PubMed as ‘Review’. We will delve deeper into the pLIRm collection in a separate work (currently in preparation), since this literature dataset was used to annotate instances of LIR-motifs which were subsequently used to train and validate computational methods presented in [19].

The distribution of publication dates for manuscripts describing experimentally verified LIR-motifs illustrates an increasing overall interest for the study of LIR-motif mediated protein-protein interactions and the progress in the field (**Figure 3**). There is an overall increasing trend in the manuscripts used to annotate LIR-motif entries in LIRcentral up to 2017 (included). The relative decline in content published after 2018 is due to the fact that (a) our massive semi-automatic analysis of full-text articles (see Materials and Methods) covered the period up to November 2020 and (b) recently, we have concentrated our efforts on developing and making available the online LIRcentral database, with a backlog of dozens of papers/LIR-motifs that are in the pipeline for inclusion in the next major release (scheduled to appear in December 2022). In any case, LIRcentral consistently covers comparable or (frequently) more papers to the other resources across the complete range from 2007 to date.

**Figure 3.**
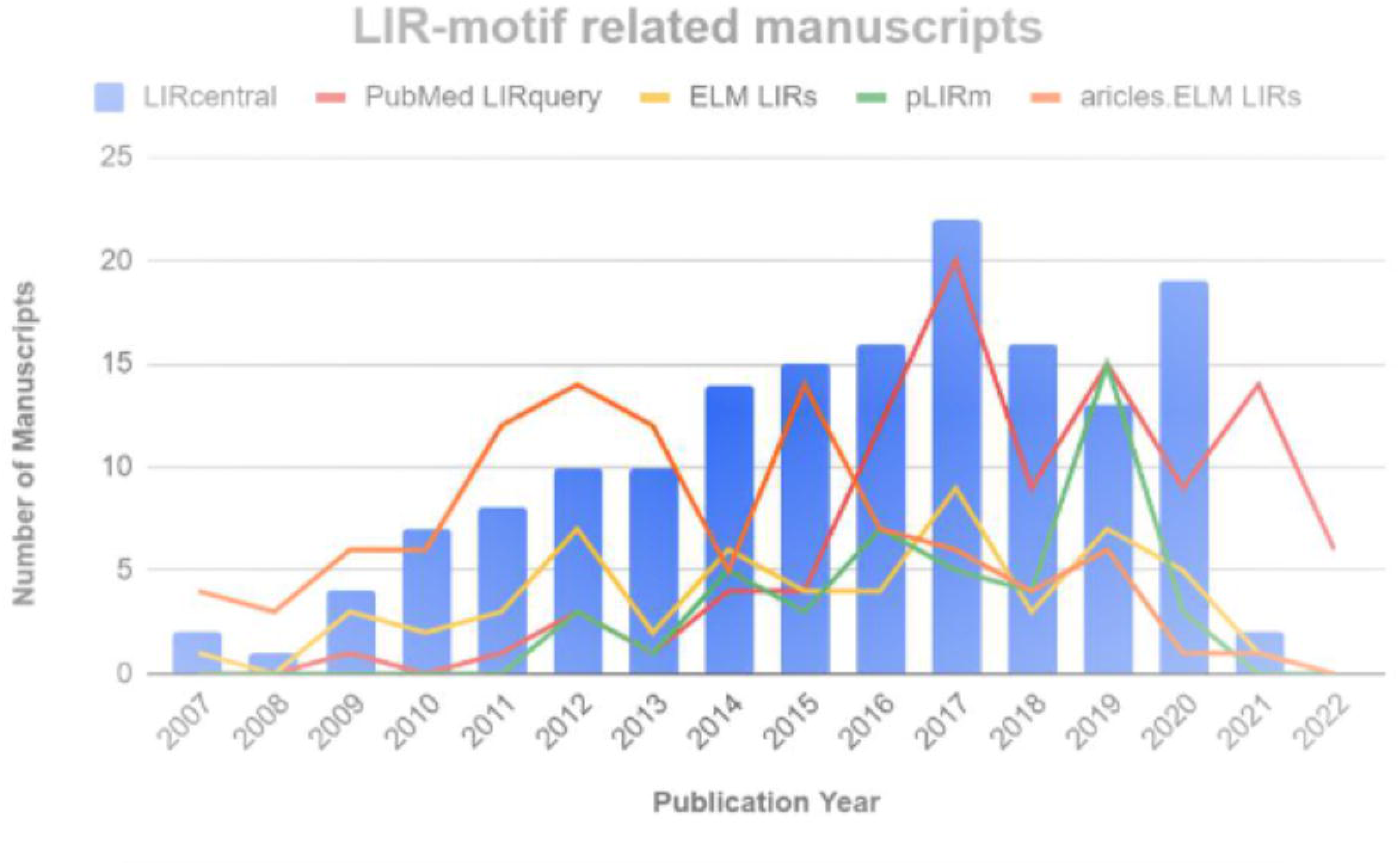
Distribution of publication dates for manuscripts describing experimentally verified LIR-motifs in the current version of LIRcentral and comparison to other literature colections of LIR-motifs.

It is worth mentioning that while developing LIRcentral, we have initially focused on covering as many LIR-motifs as possible, without spending additional resources on covering all relevant literature. That is, when a newly published paper provides additional information on the functionality of a particular motif already annotated using another primary work, we intentionally leave it aside for future curation, in order to curate papers related to motifs not currently in the database.

#### LIR-motif entries

The LIRcentral database follows a motif-centric approach, in the sense that LIRcentral entries correspond to instances of motifs (rather than protein/polypeptide entries or publications). The current LIRcentral version is composed of 292 instances of experimentally verified motifs, contained in 160 unique protein chains, originating from 22 species. Information in published manuscripts is used to identify the respective UniProt entries, and in some cases particular isoforms when this information can be deduced from the underlying curation process. LIRcentral records both functional (i.e. binding; n=201) and non-functional (n=82) instances of LIR-motifs, since negative examples can also be of great value, especially when developing prediction methods.

For the purposes of LIRcentral we identify three subclasses of binding LIR-motifs: generic (mentioned as ‘YES’) and the two special cases of (i) conditionally functional, non-canonical shuffled LIR-motifs, as observed in the human CDK5 regulatory subunit-associated protein 3 and its *Arabidopsis thaliana* homolog (UniProt: Q96JB5, Q9FG23 respectively; [28]) and (ii) accessory LIR-motifs, which function complementary to other functional (or accessory) LIR-motifs, as for example found in yeast Atg19 [29], and might play a role in defining the strength or the stoichiometry of the LC3::LIRCP interactions.

As verified but non-functional LIR-motifs (labeled as ‘NO’) we include those cases that have been unambiguously screened (e.g. via mutating the critical X_0_, X_+3_ positions of the motif) to not bind Atg8s. In several cases we can safely assume that all other peptides besides those directly shown to be functional LIR-motifs can be considered as negative examples (i.e. non-functional). Nevertheless, at this point we have decided to leave them unlabelled, since there are examples in the literature when revisiting a protein with characterized functional LIR-motifs provides updated information that is not coherent with such an inference (see for example the history of the discovery of functional LIR-motifs in yeast Atg19 [30, 29]).

In addition, there are several cases of proteins where it might be known that they interact with members of the Atg8 family in a LIR-dependent manner, yet the underlying binding motif(s) have not been characterized so far (e.g. several proteins studied in [31]). LIR-motifs predicted in such cases are designated as N/A (and can be interpreted as ‘candidate’ functional motifs), since we have not found literature that experimentally demonstrates which of the predicted motifs are indeed functional.

In addition, for all proteins with an experimentally validated LIR-motif, LIRcentral makes available all peptides matching the following regular-expression based definitions: canonical LIR-motif ([WFY]xx[VLI]), the extended LIR-motif/xLIR [16], and the five different motifs implemented in the hfAIM method [17]. Such motif instances are designated as ‘Predicted’ and are not included in the above statistics.

The vast majority of LIR-central entries comprises human LIR-motifs (168, 57.5%) followed by entries from model species such as *Saccharomyces cerevisiae* (22, 7.5%), mouse (23, 7.9%) and *Arabidopsis thaliana* (20, 6.8%). It is worth mentioning that within LIRcentral entries there are also a few LIR-motifs of non-eukaryotic origin.

### LIRcentral Access and Functionality

Users can get access to LIRcentral with any modern web browser pointed to the URL https://lircentral.eu (**Figure 4**). A simple ‘Search’ functionality facilitates entry retrieval using keyword-based queries, while the ‘Browse’ functionality offers a more advanced search interface where complex, targeted queries can be performed. Results retrieved after a successful ‘Search’ or ‘Browse’ operation are displayed in tabular form (**Figure 5**); these tables can be copied (as text in the clipboard) or exported to spreadsheet, comma-separated or PDF formatted files. Information on how to use LIRcentral is offered in a detailed online ‘Help’ page (https://lircentral.eu/help).

**Figure 4.**
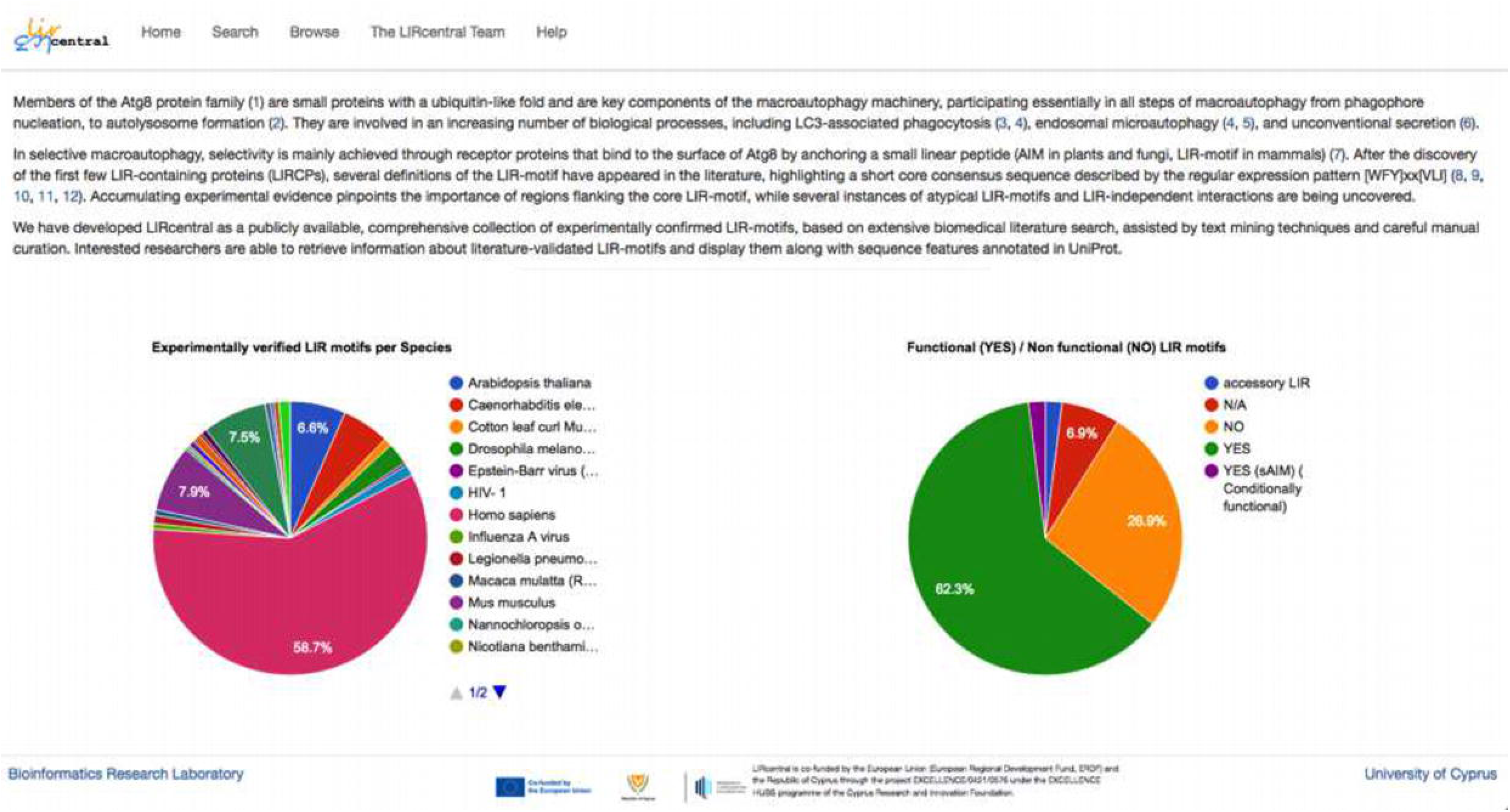
The LIRcentral landing page (‘Home’). It provides a short introduction to the purpose of LIRcentral with a brief summary of the database content and links to all LIRcentral functionalities.

**Figure 5.**
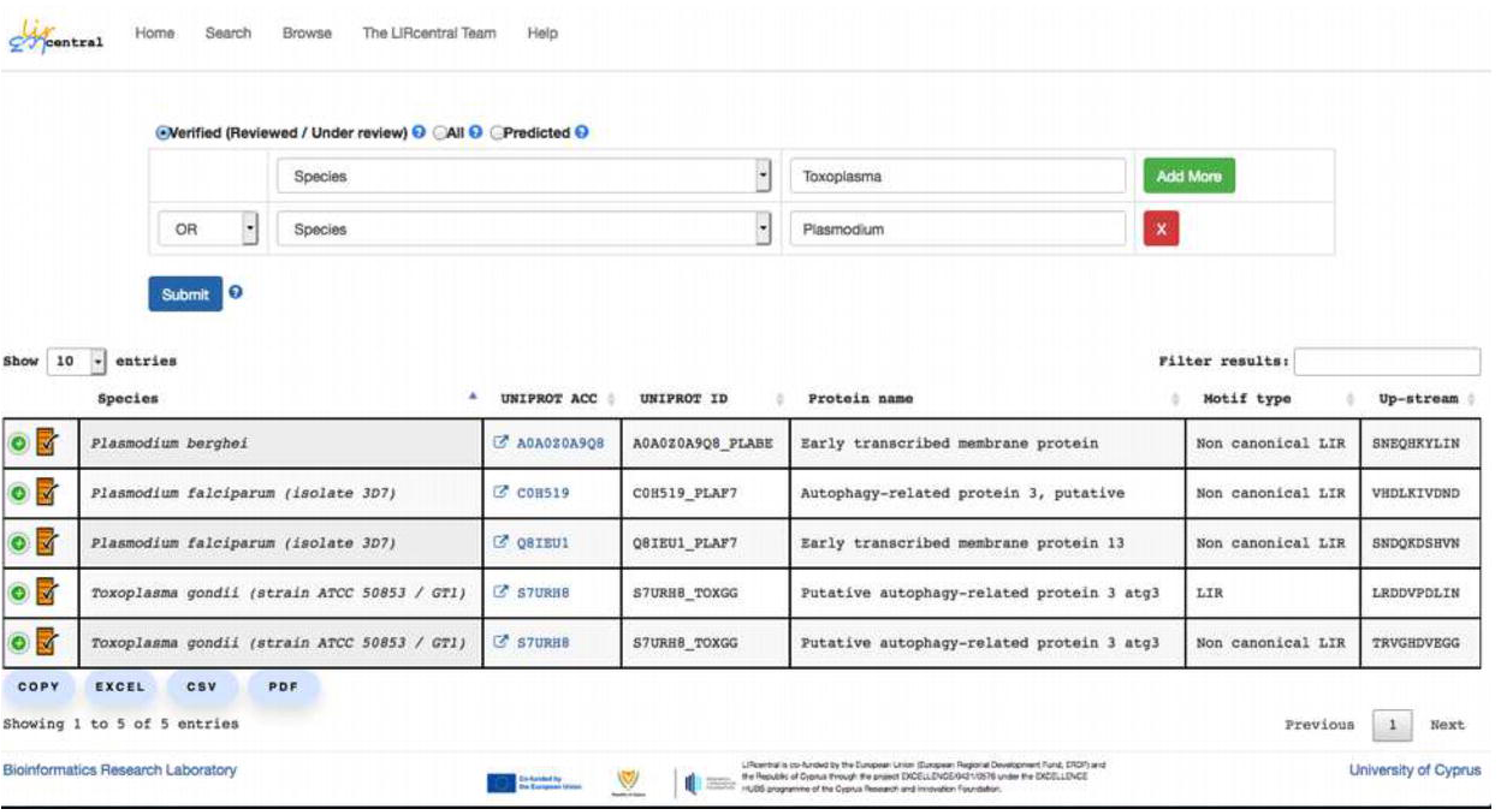
Advanced search via the ‘Browse’ functionality. Here all LIRcentral experimentally verified entries originating from species whose names match either ‘Toxoplasma’ or ‘Plasmodium’ are retrieved in a single query.

Each protein with at least one experimentally characterized LIR-motif entered in LIRcentral has its own dedicated page. There, all validated as well as predicted LIR-motifs are displayed in tabular form. A graphical illustration of LIR-motifs accompanied with features retrieved “on the fly” from the UniProt database is also provided.

### Case studies

We anticipate that LIRcentral will be used by different researchers in tackling a wide range of research questions and applications. The most straightforward use of LIRcentral will be in cases where a researcher will be interested in transferring knowledge about a LIR-motif described in the literature to a protein homolog in another species; retrieving reliable data from LIRcentral and combining them with readily available sequence analysis tools can guide the first steps for characterizing LIR-motifs in homologs. In the following, we present examples of two “larger-scale” case studies, which can greatly benefit from the capabilities offered by LIRcentral and the knowledge collected therein.

#### Case study 1 - Revealing defining sequence properties of functional LIR-motifs

Q1: Which particular residues are important for a functional LIR-motif type?
Q2: Do human LIR-motifs have distinctive sequence properties compared to motifs from other species?
Q3: What sequence properties distinguish a functional from a non-functional LIR-motif?

All the above are examples of valid (and, admittedly, interesting) biological questions that researchers working in the field might want to address (definitely, we could imagine many more questions of a similar nature!). We present a step-by-step guide on how one can use LIRcentral in combination with online tools to address the third example question above.

Ideally, it would be desirable to obtain sets of experimentally determined LIR-motifs that belong either to the functional or the non-functional class. To avoid any unwanted inhomogeneity that might occur due to different species-specific compositional biases [32, 33] we will focus on human LIR-motifs only. In addition, all instances of non-canonical LIR-motifs as well as those motifs located at the N- and C-terminal extremities of the respective LIRCPs can be excluded from the analysis.

To obtain all human LIR-motifs with experimental support, a user can navigate to the ‘Browse’ page and execute a targeted search, selecting the ‘Verified’ radio button, to restrict entries retrieved among only those which are annotated based on literature curation. By selecting the ‘Species’ option from the drop-down menu, users can enter ‘Homo sapiens’ as the target species and execute the search. The results of this query can be exported in a spreadsheet file (using the ‘EXCEL’ export button) and can be easily filtered to obtain subsets fulfilling the above criteria within minutes. For simplicity, in this case-study we remove all non-canonical human LIR-motifs and use the remaining 149 motif instances (101 functional, 48 non-functional) and use the PSSMSearch server [34] for quickly computing positional frequencies for all residue types for the core LIR-motif (**Figure 6**).

**Figure 6.**
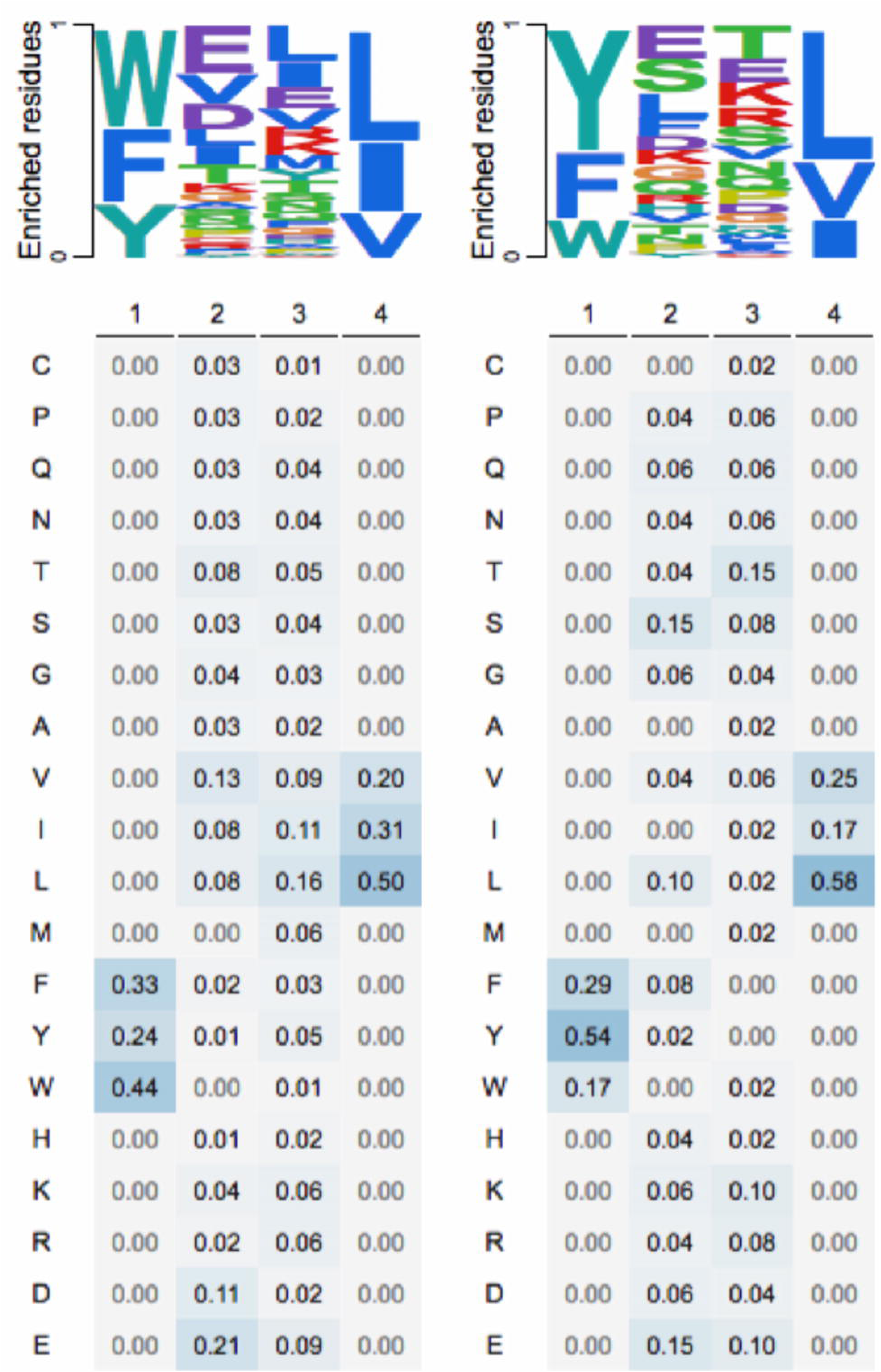
Positional frequencies of different amino acid residue types for human experimentally verified functional (left; n=101) and non-functional (right; n=48) sequences matching the canonical LIR-motif. Top: Logo-like representations; Bottom: Actual frequencies. Computations performed using PSSMsearch [34] using experimentally verified human LIR-motif data retrieved from LIRcentral.

In fact, this approach (with some methodological modifications) has already been used to develop part of a new computational scheme to predict functional LIR-motifs in *Homo sapiens* proteins (Chatzichristofi et al., in preparation). In particular, we propose the use of two PSSM models computed to represent functional and non-functional instances of human LIR-motifs along with their flanking regions, combined with the average pLDDT confidence values from AlphaFold2 predictions [35] as a proxy for the “disorderliness” of these segments [36]. These results can then be combined using a machine learning classifier for accurate prediction of human LIR-motifs.

#### Case study 2 - Constructing a Protein-Protein Interaction-driven, LIRCP network in

##### Saccharomyces cerevisiae

With the rapid advancement of -omics methods, the identification of new protein sequences is happening at an exponential rate. However, assigning biological function to a protein is still a challenging problem which often requires years of experimental work. Protein-protein interaction (PPI) networks are becoming increasingly popular, since they facilitate the enhanced description and understanding of cellular systems in physiological conditions and disease [37]. They can also aid in deciphering the biological function of proteins [38] and the prediction of protein complexes [39]. Additionally, PPI networks can provide useful insights on potential interacting partners, enabling experimental biologists to prioritize research targets. In this case study, we demonstrate how to efficiently generate a PPI network centered around LIRCPs using a list of manually curated LIRcentral entries belonging to *S. cerevisiae* (baker’s yeast).

Users can easily retrieve a list of yeast protein accessions by using the LIRcentral ‘Browse’ functionality, searching for the keyword ‘cerevisiae’ under the ‘Species’ field after selecting the ‘Verified’ radio button. This action will result in displaying a table containing all yeast entries with at least one experimentally characterized LIR-motif (21 characterized motifs, of which 18 functional in 12 different proteins). The entries can then be downloaded in any of the available formats (e.g. csv, excel). For simplicity, we downloaded results in ‘excel’ format. Using spreadsheet-like tools to further process the data, users can easily extract the list of UniProt accessions and provide them as input to tools like GeneMANIA [40] to generate a PPI network focusing on yeast LIRCPs within just a few minutes (**Figure 7**).

**Figure 7.**
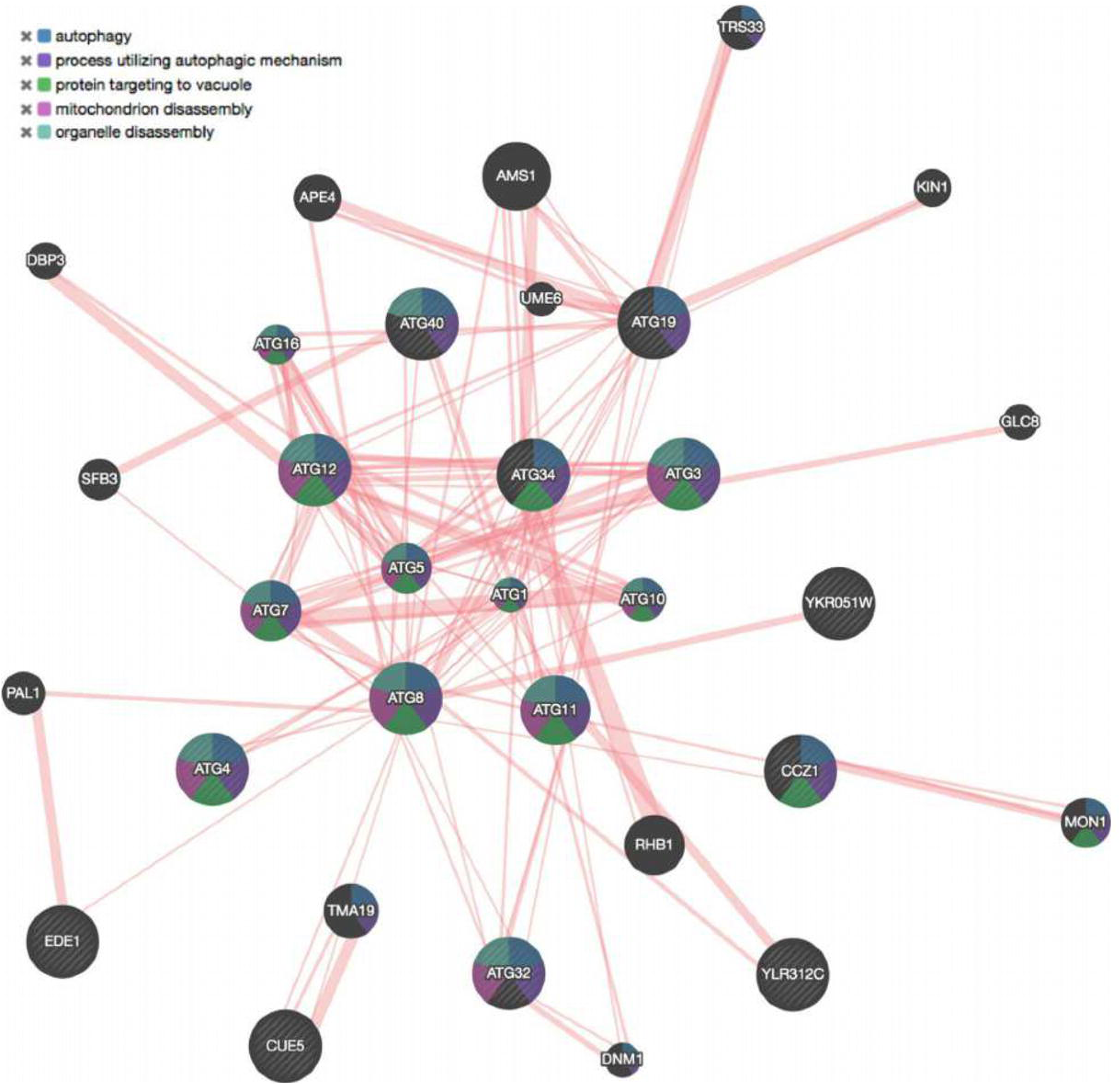
PPI network using as input a list of 12 verified yeast LIRCPs entries from LIRcentral. Proteins are represented as nodes while edges represent physical interactions between them. The input set was enhanced with the prediction of 20 additional entities based on known PPIs. Striped nodes signify input proteins and the color scheme corresponds to the top-5 gene ontology terms (FDR<10^-12^). This network was generated using the GeneMANIA web server https://genemania.org [40]: only yeast PPI data were used to infer 20 proteins related to the input dataset (all other parameters set to their default values).

## Materials and Methods

### Defining a strategy for collecting relevant literature

In order to be able to create a high quality dataset of experimentally verified LIR-motifs for inclusion in LIRcentral we had to devise a strategy for retrieving as many relevant published papers as possible to ensure high coverage. In addition, due to the significant amount of time necessary to carefully process papers for curation, a minimum number of false positives is desirable, since the limited curation resources available should not be wasted on working with irrelevant papers.

We started with an initial collection of papers, mostly retrieved by direct PubMed search (keyword-based query, i.e. using the ‘LIRquery’) or reverse search, i.e. papers citing key papers on LIR-motifs (e.g. [15]) or the software tools/database for LIR-motif prediction (e.g. [16,17,20]). Papers relevant to the description of LIR-motifs that could not be retrieved by the ‘LIRquery’ were manually inspected to identify why they are unidentifiable using direct search.

We identified several reasons why papers describing the experimental characterization of LIR-motifs do not contain a relevant mention in the title or abstract that could facilitate their automatic retrieval, the most prominent being:

1. Older papers - when the terms LIR-/AIM-motif were not yet coined or widely established (PMIDs: 17580304, 17916189, 18027972, 19021777).
2. Papers where LIR-motif characterization might not be central in the paper’s story (e.g. PMIDs: 29937374, 29277967, 28244873), where this information is ‘buried’ in the main text, in figure legends or even in supplementary data.
3. Papers mentioning LIR-motifs in a different number of (non-standardized) ways.

As an example demonstrating the latter case, we can see several different ways of expressing the concept of LIR/AIM-motifs in the literature:

‘LC3-interacting regions (LIRs)’ (PMID: 29717061)
‘LC-3 interacting regions (LIRs)’ (PMID: 30568238)

‘light chain 3 (LC3)-interacting region (LIR) motifs’ (PMID: 29937374)

‘LC3-interaction region domain’ (PMID: 29416008)

‘Atg8-family-interacting motif’ (PMID: 30451685)
‘Atg8 family-interacting motif’ (PMID: 30026233)
‘Atg8-binding sequence motif’ (PMID: 29867141)

Therefore, it became obvious that to retrieve the maximum possible number of papers for curating relevant information, we should find some simple rules based on the complete body of text included in publications, when available.

After curating a number of relevant papers, we observed that experimental evidence for the characterization of LIR-motifs is most often described in sentences containing terms with the following characteristics:

- Describe entities that refer to an Atg8 homolog (e.g. ‘LC3’, ‘Atg8’) or a LIR-motif (e.g. ‘LIR’, ‘AIM’), and
- A term (often a verb) indicating interaction (e.g. ‘interact’, ‘bind’, ‘recruit’), and
- A term suggesting a conclusion, signified by terms such as ‘confirm’, ‘identify’, ‘show’, ‘demonstrate’ etc.

In addition, there are cases where the exact sequence of the motif discovered is mentioned.

### Literature Corpus Generation and Curation

Papers describing the experimental validation of LIR-motifs are acquired using a combination of ‘bulk’ (see below) and ‘selective’ (direct and reverse PubMed search, see above) approaches. For bulk retrieval, we first download locally XML files for all papers with open access full text available in PubMedCentral matching the keyword ‘autophagy’. Such searches have been incrementally performed to cover the period up to November 2020, resulting in 78798 publications. Then, we developed a simple text analysis pipeline in the Perl programming language, implementing the rules identified to be associated with sentences describing the experimental characterization of LIR-motifs. Such snippets of text are exported in a spreadsheet-like file and highlight articles worth investigating further. A total of 5151 papers with at least one informative portion of text were further manually inspected to identify papers for adding to the curation pipeline. Even though we have not formally validated the specificity and sensitivity of this approach for identifying manuscripts describing LIR-motif::LC3 interactions, we have seen that it works well in practice (see also **Figure 2** and **Figure 3**). Also, despite a non negligible number of false positives still retained in the literature corpus to be manually examined by LIRcentral curators, the size of the corpus after this filtering is manageable.

All papers selected for curation undergo a curation process following the buddy system: at least two curators study each paper and use the information provided therein (including data supplements) to correctly identify:

(i) the species corresponding to the protein analyzed,
(ii) a UniProt identifier that unambiguously corresponds to the protein entity analyzed in each particular paper,
(iii) experimental evidence for the characterization of candidate LIR-motifs, for example mutation of key LIR-motif residues followed by binding assays against Atg8s etc.
(iv) the position of the core LIR-motif. Importantly, often the authors in the literature may report different sequence spans as the motif region. In order to have a normalized view across different proteins, what we report in LIRcetrnal is not the author-based definition but the position of the core motif, unless we are dealing with atypical motifs. One such extremely atypical case is the split LIR-motif of yeast Atg12 (UniProt: P38316), where two distal residues (Ile111, Phe185) mimic the structure of the hydrophobic and aromatic residues of a canonical LIR-motif [41].

The curations of individual “buddies” are juxtaposed and undergo final corrections before they end up in the database.

## Conclusions

We anticipate that LIRcentral will become a key resource for researchers requiring access to reliable information regarding LIRCPs and their interactions with Atg8 homologs. A small, but dedicated group of curators have made this first version of LIRcentral possible by collecting relevant papers and meticulously extracting information relevant to the experimental characterization of LIR-motifs. This information is currently impossible to be automatically extracted from papers scattered across different publication media in a non-structured manner, but it can be invaluable at the hands of both wet and computational biologists for addressing a range of research questions related to selective autophagy or other cellular processes where Atg8 family members participate. We have deliberately chosen to refrain from handling the cases of LIR-independent modes of binding to LC3 (e.g. with UIMs [42]) to maintain our focus on those interactions happening at the particular LC3::LIR-motif interface: in that sense, LIRcentral already contains several atypical LIR-motif cases.

With the advancement of our collective knowledge in the details of the underlying interactions, we expect to be able to meaningfully expand LIRcentral in future releases. For example, in mammalian species a specific subclass of LIR-motifs seems to show particular specificity towards the GABARAP clade of Atg8 homologs (GABARAP interaction motifs - GIMs [43]). This specificity seems to be reflected in the underlying sequences at the core motif but also in the flanking regions. Hopefully, we will be able to include this important piece of information within the next major release of LIRcentral. Other possible expansions with respect to the type of information recorded and made available with each LIR-motif include context-specific functionality (e.g. LIR-motif switches based on post-translational modifications) and inclusion of information related to the experimental methods used for characterizing LIR-motif functionality. In addition, we believe LIRcentral already offers a simple and intuitive interface for data retrieval. However, we are aware that certain improvements can further enhance the user experience with this resource. Some functionalities we intend to implement for providing alternative or more efficient ways to use LIRcentral include for example the possibility to support gene/protein synonyms, search functionalities based on sequence similarity.

One of the main obstacles the LIRcentral group needs to overcome is the difficulty to sustain funding for database curation. For this purpose, we have planned to develop a mechanism (including software and database components) that will enable participation of any interested parties from the research community in the literature curation process (i.e. ‘crowdsourcing’). This community effort will not only help in keeping the content up to date with the most current literature, but will also help in developing community-supported standards (e.g. how should LC3::LIR-motif interaction data be reported?) and propose solutions to difficult, ‘strategic’ problems. One example of the latter stems from the current dependence of LIRcentral from UniProt. With the foreseen possibility that an increasing number of synthetic/engineered LIR-motif containing peptides will be available in the literature (as for example in [44]), there needs to be decided whether such entries should be cataloged in LIRcentral (and how). As a first step to facilitate community-driven efforts in the context of LIRcentral, we have already partnered with the APICURON resource (https://apicuron.org/, [45]), which provides the means to monitor and expose the curation efforts of individuals, thus giving appropriate credit to all researchers who put effort in providing new or revising existing LIRcentral content.

In addition, LIRcentral participates in the preparation of the DisProt ‘Autophagy collection’ (scheduled for release in June 24, 2022), where we collaborate in the annotation of known LIR-motifs residing in experimentally verified intrinsically disordered regions [46].

Currently, an active group of curators provides content, while regular updates to LIRcentral are planned (approximately every six months). We open a call to the autophagy community for contributing in the curation effort, for example by submitting information about newly discovered (or updating information on known) LIR-motifs. We anticipate that such a ‘crowdsourcing’ of the curation process among the active autophagy research community members will (i) help towards increasing both quality and coverage of the curated data, (ii) promote the development of community-agreed ways of reporting relevant data and (iii) stimulate further research in the area of selective autophagy and engage younger researchers in the field, while at the same time it will facilitate the long-term sustainability of LIRcentral.

## Abbreviations

AIM: Atg8 interacting motif
Atg8: Autophagy-related protein 8
GABARAP: γ-aminobutyric acid receptor-associated protein
GIM: GABARAP interaction motif
LC3/MAP1LC3: Microtubule associated protein 1 light chain 3 α/β/γ
LDS: LIR docking site
LIR: LC3 interacting region
LIRCP: LIR containing protein
PMID: PubMed identifier
PPI: Protein-protein interaction
SLiM: Short Linear Motif

## Acknowledgments

The authors wish to thank former members of the Bioinformatics Research Laboratory at the University of Cyprus (especially Stelios Tsompanis) for earlier contributions to LIRcentral. We also thank the APICURON team, in particular Dr Federica Quaglia (University of Padua), for fruitful discussions and efforts for integrating LIRcentral with APICURON. We also thank Elena Papaleo for constructive comments and discussions during the LIRcentral kick-off meeting in the University of Crete (April 28-29, 2022). We are also indebted to authors of papers describing LIR-motifs who provided us access to full text when that was not available through our institutional library subscriptions.

## Funding

This work has been co-funded by the European Regional Development Fund and the Republic of Cyprus through the Cyprus Research and Innovation Foundation (Project: EXCELLENCE/0421/0576) and by the University of Cyprus [Computational approaches towards mechanistic insights and improved detection of functional LIR-motifs in selective autophagy receptors and adaptors. (idLIR)] internal grant (to V.J.P.).

## Disclosure statement

No potential conflict of interest was reported by the author(s).

## Supplementary Information

**Table S1.** The set of manuscripts returned by the ‘LIRquery’ in PubMed (accessed May 15, 2022).

**Table S2.** The 20 most recent entries retrieved from EuropePMC with the ‘LIRquery’ (accessed May 24, 2022).

## Notes

https://lircentral.eu

### Competing Interest Statement

The authors have declared no competing interest.

