## Supplementary Tables S1 and S2 for "LIRcentral: a manually curated online database of experimentally validated functional LIR-motifs"

### **Supplementary Information**

**Table S1.** The set of manuscripts returned by the ‘LIRquery’ in PubMed (accessed May 15, 2022).

**Table S2**. The 20 most recent entries retrieved from EuropePMC with the ‘LIRquery’ (accessed May 24, 2022).

**Table S1.** The set of manuscripts returned by the ‘LIRquery’ in PubMed (accessed May 15, 2022).

| **LIR-motif relevant** | **Authors** | **Title** | **Journal** | **Pub. Year** | **DOI** |
| --- | --- | --- | --- | --- | --- |
| No | Lamark T, Kirkin V, Dikic I, Johansen T. | NBR1 and p62 as cargo receptors for selective autophagy of ubiquitinated targets | Cell Cycle | 2009 | 10.4161/cc.8.13.8892 |
| Yes | Rozenknop A, Rogov VV, Rogova NY, Löhr F, Güntert P, Dikic I, Dötsch V. | Characterization of the interaction of GABARAPL-1 with the LIR motif of NBR1 | J Mol Biol | 2011 | 10.1016/j.jmb.2011.05.003 |
| Yes | Popovic D, Akutsu M, Novak I, Harper JW, Behrends C, Dikic I. | Rab GTPase-activating proteins in autophagy: regulation of endocytic and autophagy pathways by direct binding to human ATG8 modifiers | Mol Cell Biol | 2012 | 10.1128/MCB.06717-11 |
| Yes | von Muhlinen N, Akutsu M, Ravenhill BJ, Foeglein Á, Bloor S, Rutherford TJ, Freund SM, Komander D, Randow F. | LC3C, bound selectively by a noncanonical LIR motif in NDP52, is required for antibacterial autophagy | Mol Cell | 2012 | 10.1016/j.molcel.2012.08.024 |
| Yes | Alemu EA, Lamark T, Torgersen KM, Birgisdottir AB, Larsen KB, Jain A, Olsvik H, Øvervatn A, Kirkin V, Johansen T. | ATG8 family proteins act as scaffolds for assembly of the ULK complex: sequence requirements for LC3-interacting region (LIR) motifs | J Biol Chem | 2012 | 10.1074/jbc.M112.378109 |
| No | Birgisdottir ÅB, Lamark T, Johansen T. | The LIR motif - crucial for selective autophagy | J Cell Sci | 2013 | 10.1242/jcs.126128 |
| No | Wild P, McEwan DG, Dikic I. | The LC3 interactome at a glance | J Cell Sci | 2014 | 10.1242/jcs.140426 |
| No | Weissenhorn W, Fauvarque MO. | A small switch has a large effect on autophagy | Structure | 2014 | 10.1016/j.str.2013.12.006 |
| No | Kalvari I, Tsompanis S, Mulakkal NC, Osgood R, Johansen T, Nezis IP, Promponas VJ. | iLIR: A web resource for prediction of Atg8-family interacting proteins | Autophagy | 2014 | 10.4161/auto.28260 |
| Yes | Fu MM, Nirschl JJ, Holzbaur ELF. | LC3 binding to the scaffolding protein JIP1 regulates processive dynein-driven transport of autophagosomes | Dev Cell | 2014 | 10.1016/j.devcel.2014.04.015 |
| No | Popelka H, Klionsky DJ. | Analysis of the native conformation of the LIR/AIM motif in the Atg8/LC3/GABARAP-binding proteins | Autophagy | 2015 | 10.1080/15548627.2015.1111503 |
| Yes | Olsvik HL, Lamark T, Takagi K, Larsen KB, Evjen G, Øvervatn A, Mizushima T, Johansen T. | FYCO1 Contains a C-terminally Extended, LC3A/B-preferring LC3-interacting Region (LIR) Motif Required for Efficient Maturation of Autophagosomes during Basal Autophagy | J Biol Chem | 2015 | 10.1074/jbc.M115.686915 |
| Yes | Joachim J, Jefferies HB, Razi M, Frith D, Snijders AP, Chakravarty P, Judith D, Tooze SA. | Activation of ULK Kinase and Autophagy by GABARAP Trafficking from the Centrosome Is Regulated by WAC and GM130 | Mol Cell | 2015 | 10.1016/j.molcel.2015.11.018 |
| Yes | Wu F, Watanabe Y, Guo XY, Qi X, Wang P, Zhao HY, Wang Z, Fujioka Y, Zhang H, Ren JQ, Fang TC, Shen YX, Feng W, Hu JJ, Noda NN, Zhang H. | Structural Basis of the Differential Function of the Two C. elegans Atg8 Homologs, LGG-1 and LGG-2, in Autophagy | Mol Cell | 2015 | 10.1016/j.molcel.2015.11.019 |
| No | Kim BW, Kwon DH, Song HK. | Structure biology of selective autophagy receptors | BMB Rep | 2016 | 10.5483/bmbrep.2016.49.2.265 |
| No | Wu F, Wang P, Shen Y, Noda NN, Zhang H. | Small differences make a big impact: Structural insights into the differential function of the 2 Atg8 homologs in C. elegans | Autophagy | 2016 | 10.1080/15548627.2015.1137413 |
| No | Chen M, Chen Z, Wang Y, Tan Z, Zhu C, Li Y, Han Z, Chen L, Gao R, Liu L, Chen Q. | Mitophagy receptor FUNDC1 regulates mitochondrial dynamics and mitophagy | Autophagy | 2016 | 10.1080/15548627.2016.1151580 |
| No | Taniguchi K, Yamachika S, He F, Karin M. | p62/SQSTM1-Dr. Jekyll and Mr. Hyde that prevents oxidative stress but promotes liver cancer | FEBS Lett | 2016 | 10.1002/1873-3468.12301 |
| No | Jacomin AC, Samavedam S, Promponas V, Nezis IP. | iLIR database: A web resource for LIR motif-containing proteins in eukaryotes | Autophagy | 2016 | 10.1080/15548627.2016.1207016 |
| Yes | Martinez-Lopez N, Garcia-Macia M, Sahu S, Athonvarangkul D, Liebling E, Merlo P, Cecconi F, Schwartz GJ, Singh R. | Autophagy in the CNS and Periphery Coordinate Lipophagy and Lipolysis in the Brown Adipose Tissue and Liver | Cell Metab | 2016 | 10.1016/j.cmet.2015.10.008 |
| Yes | Sharifi MN, Mowers EE, Drake LE, Collier C, Chen H, Zamora M, Mui S, Macleod KF. | Autophagy Promotes Focal Adhesion Disassembly and Cell Motility of Metastatic Tumor Cells through the Direct Interaction of Paxillin with LC3 | Cell Rep | 2016 | 10.1016/j.celrep.2016.04.065 |
| Yes | Abert C, Kontaxis G, Martens S. | Accessory Interaction Motifs in the Atg19 Cargo Receptor Enable Strong Binding to the Clustered Ubiquitin-related Atg8 Protein | J Biol Chem | 2016 | 10.1074/jbc.M116.736892 |
| Yes | Mowers EE, Sharifi MN, Macleod KF. | Novel insights into how autophagy regulates tumor cell motility | Autophagy | 2016 | 10.1080/15548627.2016.1203487 |
| Yes | Noh HS, Hah YS, Zada S, Ha JH, Sim G, Hwang JS, Lai TH, Nguyen HQ, Park JY, Kim HJ, Byun JH, Hahm JR, Kang KR, Kim DR. | PEBP1, a RAF kinase inhibitory protein, negatively regulates starvation-induced autophagy by direct interaction with LC3 | Autophagy | 2016 | 10.1080/15548627.2016.1219013 |
| Yes | Kuang Y, Ma K, Zhou C, Ding P, Zhu Y, Chen Q, Xia B. | Structural basis for the phosphorylation of FUNDC1 LIR as a molecular switch of mitophagy | Autophagy | 2016 | 10.1080/15548627.2016.1238552 |
| Yes | Zhang W, Ren H, Xu C, Zhu C, Wu H, Liu D, Wang J, Liu L, Li W, Ma Q, Du L, Zheng M, Zhang C, Liu J, Chen Q. | Hypoxic mitophagy regulates mitochondrial quality and platelet activation and determines severity of I/R heart injury | Elife | 2016 | 10.7554/eLife.21407 |
| No | Fracchiolla D, Sawa-Makarska J, Martens S. | Beyond Atg8 binding: The role of AIM/LIR motifs in autophagy | Autophagy | 2017 | 10.1080/15548627.2016.1277311 |
| No | Johansen T, Birgisdottir ÅB, Huber J, Kniss A, Dötsch V, Kirkin V, Rogov VV. | Methods for Studying Interactions Between Atg8/LC3/GABARAP and LIR-Containing Proteins | Methods Enzymol | 2017 | 10.1016/bs.mie.2016.10.023 |
| No | Zheng Y, Zhang X, Chen Z. | [Research progress on mechanism of Nix-mediated mitophagy] | Zhejiang Da Xue Xue Bao Yi Xue Ban | 2017 | 10.3785/j.issn.1008-9292.2017.02.14 |
| No | Lim GG, Lim KL. | Parkin-independent mitophagy-FKBP8 takes the stage | EMBO Rep | 2017 | 10.15252/embr.201744313 |
| No | Pantoom S, Yang A, Wu YW. | Lift and cut: Anti-host autophagy mechanism of Legionella pneumophila | Autophagy | 2017 | 10.1080/15548627.2017.1327943 |
| No | Jacomin AC, Samavedam S, Charles H, Nezis IP. | iLIR@viral: A web resource for LIR motif-containing proteins in viruses | Autophagy | 2017 | 10.1080/15548627.2017.1356978 |
| No | Lee Y, Weihl CC. | Regulation of SQSTM1/p62 via UBA domain ubiquitination and its role in disease | Autophagy | 2017 | 10.1080/15548627.2017.1339845 |
| No | Joachim J, Tooze SA. | Centrosome to autophagosome signaling: Specific GABARAP regulation by centriolar satellites | Autophagy | 2017 | 10.1080/15548627.2017.1385677 |
| No | Lamark T, Svenning S, Johansen T. | Regulation of selective autophagy: the p62/SQSTM1 paradigm | Essays Biochem | 2017 | 10.1042/EBC20170035 |
| Yes | Cheng J, Liao Y, Xiao L, Wu R, Zhao S, Chen H, Hou B, Zhang X, Liang C, Xu Y, Yuan Z. | Autophagy regulates MAVS signaling activation in a phosphorylation-dependent manner in microglia | Cell Death Differ | 2017 | 10.1038/cdd.2016.121 |
| Yes | Skytte Rasmussen M, Mouilleron S, Kumar Shrestha B, Wirth M, Lee R, Bowitz Larsen K, Abudu Princely Y, O'Reilly N, Sjøttem E, Tooze SA, Lamark T, Johansen T. | ATG4B contains a C-terminal LIR motif important for binding and efficient cleavage of mammalian orthologs of yeast Atg8 | Autophagy | 2017 | 10.1080/15548627.2017.1287651 |
| Yes | Sakurai S, Tomita T, Shimizu T, Ohto U. | The crystal structure of mouse LC3B in complex with the FYCO1 LIR reveals the importance of the flanking region of the LIR motif | Acta Crystallogr F Struct Biol Commun | 2017 | 10.1107/S2053230X17001911 |
| Yes | Lee YK, Jun YW, Choi HE, Huh YH, Kaang BK, Jang DJ, Lee JA. | Development of LC3/GABARAP sensors containing a LIR and a hydrophobic domain to monitor autophagy | EMBO J | 2017 | 10.15252/embj.201696315 |
| Yes | Bhujabal Z, Birgisdottir ÅB, Sjøttem E, Brenne HB, Øvervatn A, Habisov S, Kirkin V, Lamark T, Johansen T. | FKBP8 recruits LC3A to mediate Parkin-independent mitophagy | EMBO Rep | 2017 | 10.15252/embr.201643147 |
| Yes | Yang A, Pantoom S, Wu YW. | Elucidation of the anti-autophagy mechanism of the Legionella effector RavZ using semisynthetic LC3 proteins | Elife | 2017 | 10.7554/eLife.23905 |
| Yes | Qiu Y, Zheng Y, Wu KP, Schulman BA. | Insights into links between autophagy and the ubiquitin system from the structure of LC3B bound to the LIR motif from the E3 ligase NEDD4 | Protein Sci | 2017 | 10.1002/pro.3186 |
| Yes | Kwon DH, Kim L, Kim BW, Kim JH, Roh KH, Choi EJ, Song HK. | A novel conformation of the LC3-interacting region motif revealed by the structure of a complex between LC3B and RavZ | Biochem Biophys Res Commun | 2017 | 10.1016/j.bbrc.2017.06.173 |
| Yes | Joachim J, Razi M, Judith D, Wirth M, Calamita E, Encheva V, Dynlacht BD, Snijders AP, O'Reilly N, Jefferies HBJ, Tooze SA. | Centriolar Satellites Control GABARAP Ubiquitination and GABARAP-Mediated Autophagy | Curr Biol | 2017 | 10.1016/j.cub.2017.06.021 |
| Yes | Ji MM, Lee JM, Mon H, Iiyama K, Tatsuke T, Morokuma D, Hino M, Yamashita M, Hirata K, Kusakabe T. | Lipidation of BmAtg8 is required for autophagic degradation of p62 bodies containing ubiquitinated proteins in the silkworm, Bombyx mori | Insect Biochem Mol Biol | 2017 | 10.1016/j.ibmb.2017.08.006 |
| Yes | Tusco R, Jacomin AC, Jain A, Penman BS, Larsen KB, Johansen T, Nezis IP. | Kenny mediates selective autophagic degradation of the IKK complex to control innate immune responses | Nat Commun | 2017 | 10.1038/s41467-017-01287-9 |
| No | Furuya N. | Short Overview | Methods Mol Biol | 2018 | 10.1007/7651_2017_38 |
| No | Huang X. | The potential role of HGF-MET signaling and autophagy in the war of Alectinib versus Crizotinib against ALK-positive NSCLC | J Exp Clin Cancer Res | 2018 | 10.1186/s13046-018-0707-5 |
| No | Popelka H, Klionsky DJ. | Structural basis for extremely strong binding affinity of giant ankyrins to LC3/GABARAP and its application in the inhibition of autophagy | Autophagy | 2018 | 10.1080/15548627.2018.1522884 |
| Yes | Sato M, Sato K, Tomura K, Kosako H, Sato K. | The autophagy receptor ALLO-1 and the IKKE-1 kinase control clearance of paternal mitochondria in Caenorhabditis elegans | Nat Cell Biol | 2018 | 10.1038/s41556-017-0008-9 |
| Yes | Smith MD, Harley ME, Kemp AJ, Wills J, Lee M, Arends M, von Kriegsheim A, Behrends C, Wilkinson S. | CCPG1 Is a Non-canonical Autophagy Cargo Receptor Essential for ER-Phagy and Pancreatic ER Proteostasis | Dev Cell | 2018 | 10.1016/j.devcel.2017.11.024 |
| Yes | Nüchel J, Ghatak S, Zuk AV, Illerhaus A, Mörgelin M, Schönborn K, Blumbach K, Wickström SA, Krieg T, Sengle G, Plomann M, Eckes B. | TGFB1 is secreted through an unconventional pathway dependent on the autophagic machinery and cytoskeletal regulators | Autophagy | 2018 | 10.1080/15548627.2017.1422850 |
| Yes | Zhou J, Wang Z, Wang X, Li X, Zhang Z, Fan B, Zhu C, Chen Z. | Dicot-specific ATG8-interacting ATI3 proteins interact with conserved UBAC2 proteins and play critical roles in plant stress responses | Autophagy | 2018 | 10.1080/15548627.2017.1422856 |
| Yes | Gao J, Langemeyer L, Kümmel D, Reggiori F, Ungermann C. | Molecular mechanism to target the endosomal Mon1-Ccz1 GEF complex to the pre-autophagosomal structure | Elife | 2018 | 10.7554/eLife.31145 |
| Yes | Kauffman KJ, Yu S, Jin J, Mugo B, Nguyen N, O'Brien A, Nag S, Lystad AH, Melia TJ. | Delipidation of mammalian Atg8-family proteins by each of the four ATG4 proteases | Autophagy | 2018 | 10.1080/15548627.2018.1437341 |
| No | Chu CT. | Mechanisms of selective autophagy and mitophagy: Implications for neurodegenerative diseases | Neurobiol Dis | 2019 | 10.1016/j.nbd.2018.07.015 |
| No | Turco E, Witt M, Abert C, Bock-Bierbaum T, Su MY, Trapannone R, Sztacho M, Danieli A, Shi X, Zaffagnini G, Gamper A, Schuschnig M, Fracchiolla D, Bernklau D, Romanov J, Hartl M, Hurley JH, Daumke O, Martens S. | How RB1CC1/FIP200 claws its way to autophagic engulfment of SQSTM1/p62-ubiquitin condensates | Autophagy | 2019 | 10.1080/15548627.2019.1615306 |
| No | Dutta B, Huang J, To J, Tam JP. | LIR Motif-Containing Hyperdisulfide β-Ginkgotide is Cytoprotective, Adaptogenic, and Scaffold-Ready | Molecules | 2019 | 10.3390/molecules24132417 |
| No | Mamidi AS, Ray A, Surolia N. | Structural Analysis of PfSec62-Autophagy Interacting Motifs (AIM) and PfAtg8 Interactions for Its Implications in RecovER-phagy in Plasmodium falciparum | Front Bioeng Biotechnol | 2019 | 10.3389/fbioe.2019.00240 |
| Yes | Xu Y, Zhang S, Zheng H. | The cargo receptor SQSTM1 ameliorates neurofibrillary tangle pathology and spreading through selective targeting of pathological MAPT (microtubule associated protein tau) | Autophagy | 2019 | 10.1080/15548627.2018.1532258 |
| Yes | Yang Y, Ma F, Liu Z, Su Q, Liu Y, Liu Z, Li Y. | The ER-localized Ca(2+)-binding protein calreticulin couples ER stress to autophagy by associating with microtubule-associated protein 1A/1B light chain 3 | J Biol Chem | 2019 | 10.1074/jbc.RA118.005166 |
| Yes | Wang ZT, Lu MH, Zhang Y, Ji WL, Lei L, Wang W, Fang LP, Wang LW, Yu F, Wang J, Li ZY, Wang JR, Wang TH, Dou F, Wang QW, Wang XL, Li S, Ma QH, Xu RX. | Disrupted-in-schizophrenia-1 protects synaptic plasticity in a transgenic mouse model of Alzheimer's disease as a mitophagy receptor | Aging Cell | 2019 | 10.1111/acel.12860 |
| Yes | Padman BS, Nguyen TN, Uoselis L, Skulsuppaisarn M, Nguyen LK, Lazarou M. | LC3/GABARAPs drive ubiquitin-independent recruitment of Optineurin and NDP52 to amplify mitophagy | Nat Commun | 2019 | 10.1038/s41467-019-08335-6 |
| Yes | Birgisdottir ÅB, Mouilleron S, Bhujabal Z, Wirth M, Sjøttem E, Evjen G, Zhang W, Lee R, O'Reilly N, Tooze SA, Lamark T, Johansen T. | Members of the autophagy class III phosphatidylinositol 3-kinase complex I interact with GABARAP and GABARAPL1 via LIR motifs | Autophagy | 2019 | 10.1080/15548627.2019.1581009 |
| Yes | Zhang Y, Yao Y, Qiu X, Wang G, Hu Z, Chen S, Wu Z, Yuan N, Gao H, Wang J, Song H, Girardin SE, Qian Y. | Listeria hijacks host mitophagy through a novel mitophagy receptor to evade killing | Nat Immunol | 2019 | 10.1038/s41590-019-0324-2 |
| Yes | Kang HM, Noh KH, Chang TK, Park D, Cho HS, Lim JH, Jung CR. | Ubiquitination of MAP1LC3B by pVHL is associated with autophagy and cell death in renal cell carcinoma | Cell Death Dis | 2019 | 10.1038/s41419-019-1520-6 |
| Yes | Jeon P, Park JH, Jun YW, Lee YK, Jang DJ, Lee JA. | Development of GABARAP family protein-sensitive LIR-based probes for neuronal autophagy | Mol Brain | 2019 | 10.1186/s13041-019-0458-z |
| Yes | Chino H, Hatta T, Natsume T, Mizushima N. | Intrinsically Disordered Protein TEX264 Mediates ER-phagy | Mol Cell | 2019 | 10.1016/j.molcel.2019.03.033 |
| Yes | Wirth M, Zhang W, Razi M, Nyoni L, Joshi D, O'Reilly N, Johansen T, Tooze SA, Mouilleron S. | Molecular determinants regulating selective binding of autophagy adapters and receptors to ATG8 proteins | Nat Commun | 2019 | 10.1038/s41467-019-10059-6 |
| Yes | Holdgaard SG, Cianfanelli V, Pupo E, Lambrughi M, Lubas M, Nielsen JC, Eibes S, Maiani E, Harder LM, Wesch N, Foged MM, Maeda K, Nazio F, de la Ballina LR, Dötsch V, Brech A, Frankel LB, Jäättelä M, Locatelli F, Barisic M, Andersen JS, Bekker-Jensen S, Lund AH, Rogov VV, Papaleo E, Lanzetti L, De Zio D, Cecconi F. | Selective autophagy maintains centrosome integrity and accurate mitosis by turnover of centriolar satellites | Nat Commun | 2019 | 10.1038/s41467-019-12094-9 |
| No | Holdgaard SG, Cianfanelli V, Cecconi F. | Cloud hunting: doryphagy, a form of selective autophagy that degrades centriolar satellites | Autophagy | 2020 | 10.1080/15548627.2019.1703356 |
| No | Popelka H, Klionsky DJ. | Molecular dynamics simulations reveal how the reticulon-homology domain of the autophagy receptor RETREG1/FAM134B remodels membranes for efficient selective reticulophagy | Autophagy | 2020 | 10.1080/15548627.2020.1719725 |
| No | Sora V, Kumar M, Maiani E, Lambrughi M, Tiberti M, Papaleo E. | Structure and Dynamics in the ATG8 Family From Experimental to Computational Techniques | Front Cell Dev Biol | 2020 | 10.3389/fcell.2020.00420 |
| No | Jacomin AC, Petridi S, Di Monaco M, Nezis IP. | A nuclear role for Atg8-family proteins | Autophagy | 2020 | 10.1080/15548627.2020.1794356 |
| No | Wesch N, Kirkin V, Rogov VV. | Atg8-Family Proteins-Structural Features and Molecular Interactions in Autophagy and Beyond | Cells | 2020 | 10.3390/cells9092008 |
| Yes | Huber J, Obata M, Gruber J, Akutsu M, Löhr F, Rogova N, Güntert P, Dikic I, Kirkin V, Komatsu M, Dötsch V, Rogov VV. | An atypical LIR motif within UBA5 (ubiquitin like modifier activating enzyme 5) interacts with GABARAP proteins and mediates membrane localization of UBA5 | Autophagy | 2020 | 10.1080/15548627.2019.1606637 |
| Yes | Catarino S, Ribeiro-Rodrigues TM, Sá Ferreira R, Ramalho J, Abert C, Martens S, Girão H. | A Conserved LIR Motif in Connexins Mediates Ubiquitin-Independent Binding to LC3/GABARAP Proteins | Cells | 2020 | 10.3390/cells9040902 |
| Yes | Jacomin AC, Petridi S, Di Monaco M, Bhujabal Z, Jain A, Mulakkal NC, Palara A, Powell EL, Chung B, Zampronio C, Jones A, Cameron A, Johansen T, Nezis IP. | Regulation of Expression of Autophagy Genes by Atg8a-Interacting Partners Sequoia, YL-1, and Sir2 in Drosophila | Cell Rep | 2020 | 10.1016/j.celrep.2020.107695 |
| Yes | Li Y, Cheng X, Li M, Wang Y, Fu T, Zhou Z, Wang Y, Gong X, Xu X, Liu J, Pan L. | Decoding three distinct states of the Syntaxin17 SNARE motif in mediating autophagosome-lysosome fusion | Proc Natl Acad Sci U S A | 2020 | 10.1073/pnas.2006997117 |
| No | Di Rita A, Strappazzon F. | A protective variant of the autophagy receptor CALCOCO2/NDP52 in Multiple Sclerosis (MS) | Autophagy | 2021 | 10.1080/15548627.2021.1924969 |
| No | Zhang W, Han Z, Xue Y, Jia D. | iCAL: a new pipeline to investigate autophagy selectivity and cancer | Autophagy | 2021 | 10.1080/15548627.2021.1939972 |
| No | Abudu YP, Mouilleron S, Tooze SA, Lamark T, Johansen T. | SAMM50 is a receptor for basal piecemeal mitophagy and acts with SQSTM1/p62 in OXPHOS-induced mitophagy | Autophagy | 2021 | 10.1080/15548627.2021.1953846 |
| No | Tóth D, Horváth GV, Juhász G. | The interplay between pathogens and Atg8 family proteins: thousand-faced interactions | FEBS Open Bio | 2021 | 10.1002/2211-5463.13318 |
| Yes | Wang R, Zhu Y, Ren C, Yang S, Tian S, Chen H, Jin M, Zhou H. | Influenza A virus protein PB1-F2 impairs innate immunity by inducing mitophagy | Autophagy | 2021 | 10.1080/15548627.2020.1725375 |
| Yes | Fraiberg M, Tamim-Yecheskel BC, Kokabi K, Subic N, Heimer G, Eck F, Nalbach K, Behrends C, Ben-Zeev B, Shatz O, Elazar Z. | Lysosomal targeting of autophagosomes by the TECPR domain of TECPR2 | Autophagy | 2021 | 10.1080/15548627.2020.1852727 |
| Yes | Shu L, Hu C, Xu M, Yu J, He H, Lin J, Sha H, Lu B, Engelender S, Guan M, Song Z. | ATAD3B is a mitophagy receptor mediating clearance of oxidative stress-induced damaged mitochondrial DNA | EMBO J | 2021 | 10.15252/embj.2020106283 |
| Yes | Chang HC, Tao RN, Tan CT, Wu YJ, Bay BH, Yu VC. | The BAX-binding protein MOAP1 associates with LC3 and promotes closure of the phagophore | Autophagy | 2021 | 10.1080/15548627.2021.1896157 |
| Yes | Wirth M, Mouilleron S, Zhang W, Sjøttem E, Princely Abudu Y, Jain A, Lauritz Olsvik H, Bruun JA, Razi M, Jefferies HBJ, Lee R, Joshi D, O'Reilly N, Johansen T, Tooze SA. | Phosphorylation of the LIR Domain of SCOC Modulates ATG8 Binding Affinity and Specificity | J Mol Biol | 2021 | 10.1016/j.jmb.2021.166987 |
| Yes | Carinci M, Testa B, Bordi M, Milletti G, Bonora M, Antonucci L, Ferraina C, Carro M, Kumar M, Ceglie D, Eck F, Nardacci R, le Guerroué F, Petrini S, Soriano ME, Caruana I, Doria V, Manifava M, Peron C, Lambrughi M, Tiranti V, Behrends C, Papaleo E, Pinton P, Giorgi C, Ktistakis NT, Locatelli F, Nazio F, Cecconi F. | TFG binds LC3C to regulate ULK1 localization and autophagosome formation | EMBO J | 2021 | 10.15252/embj.2019103563 |
| Yes | Abudu YP, Shrestha BK, Zhang W, Palara A, Brenne HB, Larsen KB, Wolfson DL, Dumitriu G, Øie CI, Ahluwalia BS, Levy G, Behrends C, Tooze SA, Mouilleron S, Lamark T, Johansen T. | SAMM50 acts with p62 in piecemeal basal- and OXPHOS-induced mitophagy of SAM and MICOS components | J Cell Biol | 2021 | 10.1083/jcb.202009092 |
| Yes | Liu M, Zhang W, Li M, Feng J, Kuang W, Chen X, Yang F, Sun Q, Xu Z, Hua J, Yang C, Liu W, Shu Q, Yang Y, Zhou T, Xie S. | NudCL2 is an autophagy receptor that mediates selective autophagic degradation of CP110 at mother centrioles to promote ciliogenesis | Cell Res | 2021 | 10.1038/s41422-021-00560-3 |
| Yes | Poole LP, Bock-Hughes A, Berardi DE, Macleod KF. | ULK1 promotes mitophagy via phosphorylation and stabilization of BNIP3 | Sci Rep | 2021 | 10.1038/s41598-021-00170-4 |
| Yes | Ordureau A, Kraus F, Zhang J, An H, Park S, Ahfeldt T, Paulo JA, Harper JW. | Temporal proteomics during neurogenesis reveals large-scale proteome and organelle remodeling via selective autophagy | Mol Cell | 2021 | 10.1016/j.molcel.2021.10.001 |
| No | Popelka H, Klionsky DJ. | The RB1CC1 Claw-binding motif: a new piece in the puzzle of autophagy regulation | Autophagy | 2022 | 10.1080/15548627.2022.2029234 |
| No | Tsapras P, Nezis IP. | A yeast two-hybrid screening identifies novel Atg8a interactors in Drosophila | Autophagy | 2022 | 10.1080/15548627.2022.2045535 |
| No | Zhang H, Sun H, Zhang W, Xu Y, Geng D. | Identification of Key Genes and Potential Mechanisms Based on the Autophagy Regulatory Network in Osteoclasts Using a Murine Osteoarthritis Model | J Inflamm Res | 2022 | 10.2147/JIR.S354824 |
| Yes | Zhao J, Li Z, Li J. | The crystal structure of the FAM134B-GABARAP complex provides mechanistic insights into the selective binding of FAM134 to the GABARAP subfamily | FEBS Open Bio | 2022 | 10.1002/2211-5463.13340 |
| Yes | Qin X, Wang R, Xu H, Tu L, Chen H, Li H, Liu N, Wang J, Li S, Yin F, Xu N, Li Z. | Identification of an autoinhibitory, mitophagy-inducing peptide derived from the transmembrane domain of USP30 | Autophagy | 2022 | 10.1080/15548627.2021.2022360 |
| Yes | B B, Zeng Z, Zhou C, Lian G, Guo F, Wang J, Han N, Zhu M, Bian H. | Identification of New ATG8s-Binding Proteins with Canonical LC3-Interacting Region in Autophagosomes of Barley Callus | Plant Cell Physiol | 2022 | 10.1093/pcp/pcac015 |

The first column (‘LIR-motif relevant’) marks papers describing primary experimental data for the characterization of LIR-motifs:

‘No’ (shown with blue background) signifies papers which do not present primary data on the experimental characterization of LIR-motifs. ‘Yes’ (shown with green background) signifies papers which present primary data on the experimental characterization of at least one LIR-motif.

**Table S2**. The 20 most recent entries retrieved from EuropePMC with the ‘LIRquery’ (accessed May 24, 2022).

| **LIR-motif**  **relevant** | **Authors** | **Title** | **Journal** | **Pub. Date** | **DOI** |
| --- | --- | --- | --- | --- | --- |
| No | Quinet, Grégoire; Génin, Pierre; Ozturk, Oznur; Belgareh-Touzé, Naima; Courtot, Lilas; Legouis, Renaud; Weil, Robert; Cohen, Mickael M; Rodriguez, Manuel S | Exploring selective autophagy events in multiple biologic models using LC3-interacting regions (LIR)-based molecular traps. | Sci Rep | 2022-05-10 | 10.1038/s41598-022-11417-z |
| No | Li, Anqi; Gao, Meng; Liu, Bilin; Qin, Yuan; Chen, Lei; Liu, Hanyu; Wu, Huayan; Gong, Guohua | Mitochondrial autophagy: molecular mechanisms and implications for cardiovascular disease. | Cell Death Dis | 2022-05-09 | 10.1038/s41419-022-04906-6 |
| No | Fassi, Enrico Mario Alessandro; Garofalo, Mariangela; Sgrignani, Jacopo; Dei Cas, Michele; Mori, Matteo; Roda, Gabriella; Cavalli, Andrea; Grazioso, Giovanni | Focused Design of Novel Cyclic Peptides Endowed with GABARAP-Inhibiting Activity. | Int J Mol Sci | 2022-05-03 | 10.3390/ijms23095070 |
| No | Lu, Yue; He, Ping; Zhang, Yuxuan; Ren, Yongwen; Zhang, Leiliang | The emerging roles of retromer and sorting nexins in the life cycle of viruses. | Virol Sin | 2022-05-02 | 10.1016/j.virs.2022.04.014 |
| N/A - duplicate entry | Fassi, Enrico Mario Alessandro; Garofalo, Mariangela; Sgrignani, Jacopo; Dei Cas, Michele; Mori, Matteo; Roda, Gabriella; Cavalli, Andrea; Grazioso, Giovanni | Focused Design of Novel Cyclic Peptides Endowed with GABARAP-Inhibiting Activity | Int J Mol Sci | 2022-05-01 |  |
| Yes | Moyzis, Alexandra G.; Lally, Navraj S.; Liang, Wenjing; Najor, Rita H.; Gustafsson, Åsa B. | Mcl-1 Differentially Regulates Autophagy in Response to Changes in Energy Status and Mitochondrial Damage | Cells | 2022-04-27 |  |
| Yes | Shi, Xiao Chen; Xia, Bo; Zhang, Jian Feng; Zhang, Rui Xin; Zhang, Dan Yang; Liu, Huan; Xie, Bao Cai; Wang, Yong Liang; Wu, Jiang Wei | Optineurin promotes myogenesis during muscle regeneration in mice by autophagic degradation of GSK3β. | PLoS Biol | 2022-04-27 | 10.1371/journal.pbio.3001619 |
| No | Vianello, Caterina; Cocetta, Veronica; Catanzaro, Daniela; Dorn, Gerald W; De Milito, Angelo; Rizzolio, Flavio; Canzonieri, Vincenzo; Cecchin, Erika; Roncato, Rossana; Toffoli, Giuseppe; Quagliariello, Vincenzo; Di Mauro, Annabella; Losito, Simona; Maurea, Nicola; Cono, Scaffa; Sales, Gabriele; Scorrano, Luca; Giacomello, Marta; Montopoli, Monica | Cisplatin resistance can be curtailed by blunting Bnip3-mediated mitochondrial autophagy. | Cell Death Dis | 2022-04-22 | 10.1038/s41419-022-04741-9 |
| No | Huh, Sung Un | Evolutionary Diversity and Function of Metacaspases in Plants: Similar to but Not Caspases | Int J Mol Sci | 2022-04-21 |  |
| No | Chipurupalli, Sandhya; Ganesan, Raja; Martini, Giulia; Mele, Luigi; Reggio, Alessio; Esposito, Marianna; Kannan, Elango; Namasivayam, Vigneshwaran; Grumati, Paolo; Desiderio, Vincenzo; Robinson, Nirmal | Cancer cells adapt FAM134B/BiP mediated ER-phagy to survive hypoxic stress. | Cell Death Dis | 2022-04-18 | 10.1038/s41419-022-04813-w |
| No | Park, Na Yeon; Jo, Doo Sin; Cho, Dong-Hyung | Post-Translational Modifications of ATG4B in the Regulation of Autophagy. | Cells | 2022-04-13 | 10.3390/cells11081330 |
| No | Zhang, Haifeng; Sun, Houyi; Zhang, Wei; Xu, Yaozeng; Geng, Dechun | Identification of Key Genes and Potential Mechanisms Based on the Autophagy Regulatory Network in Osteoclasts Using a Murine Osteoarthritis Model. | J Inflamm Res | 2022-04-12 | 10.2147/jir.s354824 |
| Yes | Peng, Jialing; Pan, Jingrui; Wang, Hongxuan; Mo, Jingjing; Lan, Lihuan; Peng, Ying | Morphine-induced microglial immunosuppression via activation of insufficient mitophagy regulated by NLRX1. | J Neuroinflammation | 2022-04-12 | 10.1186/s12974-022-02453-7 |
| No | Liu, Kun; Zhao, Qian; Sun, Hongyan; Liu, Lei; Wang, Chaoqun; Li, Zheng; Xu, Youqing; Wang, Liang; Zhang, Lin; Zhang, Honghai; Chen, Quan; Zhao, Tongbiao | BNIP3 (BCL2 interacting protein 3) regulates pluripotency by modulating mitochondrial homeostasis via mitophagy. | Cell Death Dis | 2022-04-11 | 10.1038/s41419-022-04795-9 |
| No | Li, Qin; Liu, Yinghai; Huang, Qingqing; Yi, Xiaobo; Qin, Fuen; Zhong, Zuling; Lin, Lu; Yang, Haihong; Gong, Gu; Wu, Wei | Hypoxia Acclimation Protects against Heart Failure Postacute Myocardial Infarction via Fundc1-Mediated Mitophagy. | Oxid Med Cell Longev | 2022-04-05 | 10.1155/2022/8192552 |
| No | Zhao, Lin; Zhao, Jia; Zhong, Kunhong; Tong, Aiping; Jia, Da | Targeted protein degradation: mechanisms, strategies and application. | Signal Transduct Target Ther | 2022-04-04 | 10.1038/s41392-022-00966-4 |
| No | Guan, Xinjie; Iyaswamy, Ashok; Sreenivasmurthy, Sravan Gopalkrishnashetty; Su, Chengfu; Zhu, Zhou; Liu, Jia; Kan, Yuxuan; Cheung, King-Ho; Lu, Jiahong; Tan, Jieqiong; Li, Min | Mechanistic Insights into Selective Autophagy Subtypes in Alzheimer's Disease. | Int J Mol Sci | 2022-03-25 | 10.3390/ijms23073609 |
| No | Xu, Dan-Dan; Du, Li-Lin | Fission Yeast Autophagy Machinery. | Cells | 2022-03-24 | 10.3390/cells11071086 |
| No | Jetto, Cuckoo Teresa; Nambiar, Akshaya; Manjithaya, Ravi | Mitophagy and Neurodegeneration: Between the Knowns and the Unknowns. | Front Cell Dev Biol | 2022-03-22 | 10.3389/fcell.2022.837337 |
| No | Zhou, Huimin; Wang, Kexin; Wang, Mengyan; Zhao, Wenxia; Zhang, Conghui; Cai, Meilian; Qiu, Yuhan; Zhang, Tianshu; Shao, Rongguang; Zhao, Wuli | ER-phagy in the Occurrence and Development of Cancer. | Biomedicines | 2022-03-18 | 10.3390/biomedicines10030707 |

Columns as in Table S1. One duplicate entry was retrieved and marked as ‘N/A - duplicate entry’.
